# Abundance versus presence–absence in extinction debt: Time-lagged effects of land use change on salamander populations

**DOI:** 10.64898/2026.02.09.704953

**Authors:** Yusuke Tamada, Nozomi Ohuchi, Kyosuke Shizukuda

**Author notes:** Forestry Research Institute, Hokkaido Research Organization, Higashiyama, Koshunaicho, Bibai, Hokkaido, 079-0198, Japan (Yusuke Tamada). **Declaration of authorship** Corresponding author: Yusuke Tamada.

## Abstract

Habitat loss and fragmentation have caused global biodiversity loss. It is important to understand the impact of past land use on biodiversity, not only in the present, but also to inform appropriate land-use strategies. Most studies on extinction debt, the concept that past land use drives present organism distribution, focus on species richness. However, when focusing on individual species, species richness is based on presence–absence data and does not reflect abundance. Therefore, extinction debt may not be clearly evident in presence–absence data. However, few studies have compared the effects of past land use on organism distribution using both abundance and presence–absence data. In this study, we compared the effects of past land use on the abundance and presence–absence of the ezo salamander *Hynobius retardatus*. We surveyed the aquatic areas of Ezo salamander egg sacs along a survey route in northern Japan. We measured the percentages of past and present forest areas, the physical environment, water quality, and waterbody type. The number of egg sacs was significantly associated with the percentage of past forest area, physical environment, water quality, and waterbody type. The presence or absence of egg sacs was significantly associated with the physical environment and water quality. Our results suggest that the effects of past land use on organism distribution are more evident in abundance data than in presence–absence data. Researchers and land managers may need to consider the time lag before extinction based on abundance data to assess extinction risk accurately.

## Introduction

Habitat loss and fragmentation due to urbanization, conversion to agricultural land, and forestry have caused biodiversity loss worldwide (Tilman et al. 2001; Grimm et al. 2008; Newbold et al. 2015). Habitat loss and fragmentation reduce populations through edge effects, loss of connectivity, and reduced genetic diversity (Hanski 2011; Exposito-Alonso et al. 2022). However, recent studies have shown a general trend of increasing forest area in developed countries in the Northern Hemisphere, which has been attributed to two main factors: the abandonment of agricultural land in the context of population decline and the implementation of forest conservation measures (Queiroz et al. 2014; Bryan et al. 2018; Winkler et al. 2021). Although these trends have raised hope for habitat restoration and rewilding (Zhang et al. 2010; Navarro and Pereira 2012), forest expansion does not necessarily imply biodiversity recovery. The legacy of historical land use can strongly influence biodiversity; therefore, failing to consider these effects can lead to misinterpretation of extinction risk and recovery dynamics (Tilman et al. 1994; Hanski and Ovaskainen 2002). Therefore, it is important to understand the effects of both present and past land use on biodiversity and consider appropriate land use strategies.

There is a time lag between landscape changes and their impact on organism distribution, and that past land use determines present organism distribution (Kuussaari et al. 2009; Jackson and Sax 2010). This phenomenon, known as ‘extinction debt,’ has been particularly reported in long-lived species and species that reproduce by cloning (Kuussaari et al. 2009). However, most of this research has focused on plants and invertebrates in temperate regions, leaving a knowledge gap regarding vertebrates (Figueiredo et al. 2019). Furthermore, most studies on extinction debt have focused on species richness (Hylander and Ehrlén 2013). The extinction debt related to species richness results from a time lag in the extinction of each species. However, each species responds to landscape changes in its own way. Therefore, it is important to understand these mechanisms at the species level to elucidate the underlying mechanisms of the extinction debt. Furthermore, when focusing on species richness, the data comprises presence–absence data, which do not reflect abundance. Species extinction does not occur until the abundance reaches zero, and in the case of presence–absence data, extinction debt may not be clearly reflected. Indeed, it has been reported that the impact of present land use on organism distribution is more pronounced when individual data are used than when presence–absence data are used (Cushman and McGarigal 2004; Thornton et al. 2011). However, no studies have compared the effects of past land use on the presence–absence, and abundance of species.

Amphibians are among the world’s most endangered animal species, and are affected by factors such as habitat loss and fragmentation, climate change, and infectious diseases (Pounds et al. 2006; Cordier et al. 2021; Guirguis et al. 2023). Salamanders play an important ecological role in stabilizing food webs and nutrient cycles by acting as intermediate predators between aquatic and terrestrial environments at different life stages (Davic and Welsh 2004; Best and Welsh 2014). However, 39% of frogs, 16% of caecilians, and 60% of salamanders are threatened with extinction (Re:wild et al. 2023), making salamanders among the most endangered taxa worldwide.

The Japanese archipelago has high salamander diversity and endemism, with 56 species belonging to 3 families and 6 genera confirmed to date (Herpetological Society of Japan 2025). Among them, the Ezo salamander *Hynobius retardatus*, which is endemic to Hokkaido, northern Japan, lays eggs in ponds, puddles, and ditches in the spring, after which adults move to nearby forests to inhabit (Sato 2013a; Tokuda 2022). This species has a long lifespan and has survived for over 10 years in the wild (Ueta and Sato 2013). Additionally, this species is not very mobile and is dependent on the forest landscape (Fuyuki et al. 2014; Tran et al. 2021). Therefore, past changes in the forest landscape may have affected the distribution of Ezo salamanders.

In this study, we hypothesized that the effects of past land use on Ezo salamanders would differ depending on their presence – absence or abundance. Specifically, when presence–absence data are used as the response variable, the effects of past land use may not be significant, whereas when abundance data are used, they may be significant. Therefore, we aimed to compare the effects of past land use on organism distribution in terms of presence–absence and abundance, and to evaluate the effects of past forest area on the distribution of Ezo salamanders at 50 study sites across five regions in Hokkaido, northern Japan.

## Materials and methods

### Study area

The study area covered five regions in Hokkaido, northern Japan: Sapporo, Ebetsu, Ashibetsu, Asahikawa, and Nayoro (Table 1 and Figure 1; 42°02’N–44°00’N, 141°00’E–142°27’E). We established survey routes ranging from 3.5 to 12.5 km in each region (Table 1). The annual average temperature ranged from 5.8 to 9.2 °C, and the annual average precipitation ranged from 965.0 to 1146.1 mm. The land cover in each region mainly consists of deciduous broad-leaved forests dominated by *Acer pictum*, *Quercus crispula*, and *Betula platyphylla*. Urban areas, croplands, and paddy fields are distributed throughout the region. Sapporo, the central city of Hokkaido, has a population of approximately 1.97 million. The survey route was set in the foothills adjacent to the urban area. The Ebetsu survey route is located in a hill forest surrounded by urban areas and agricultural land. During World War II, large areas of natural forests in this region were cut to produce military equipment (Koshika and Nigi, 1998). The Ashibetsu survey route currently consists mainly of deciduous broadleaved forests; however, the surrounding area was used for coal mining and mine railways until the 1960s. The areas surrounding the survey routes in Asahikawa and Nayoro have been used as rice paddies since the 1960s.

**Fig. 1.**
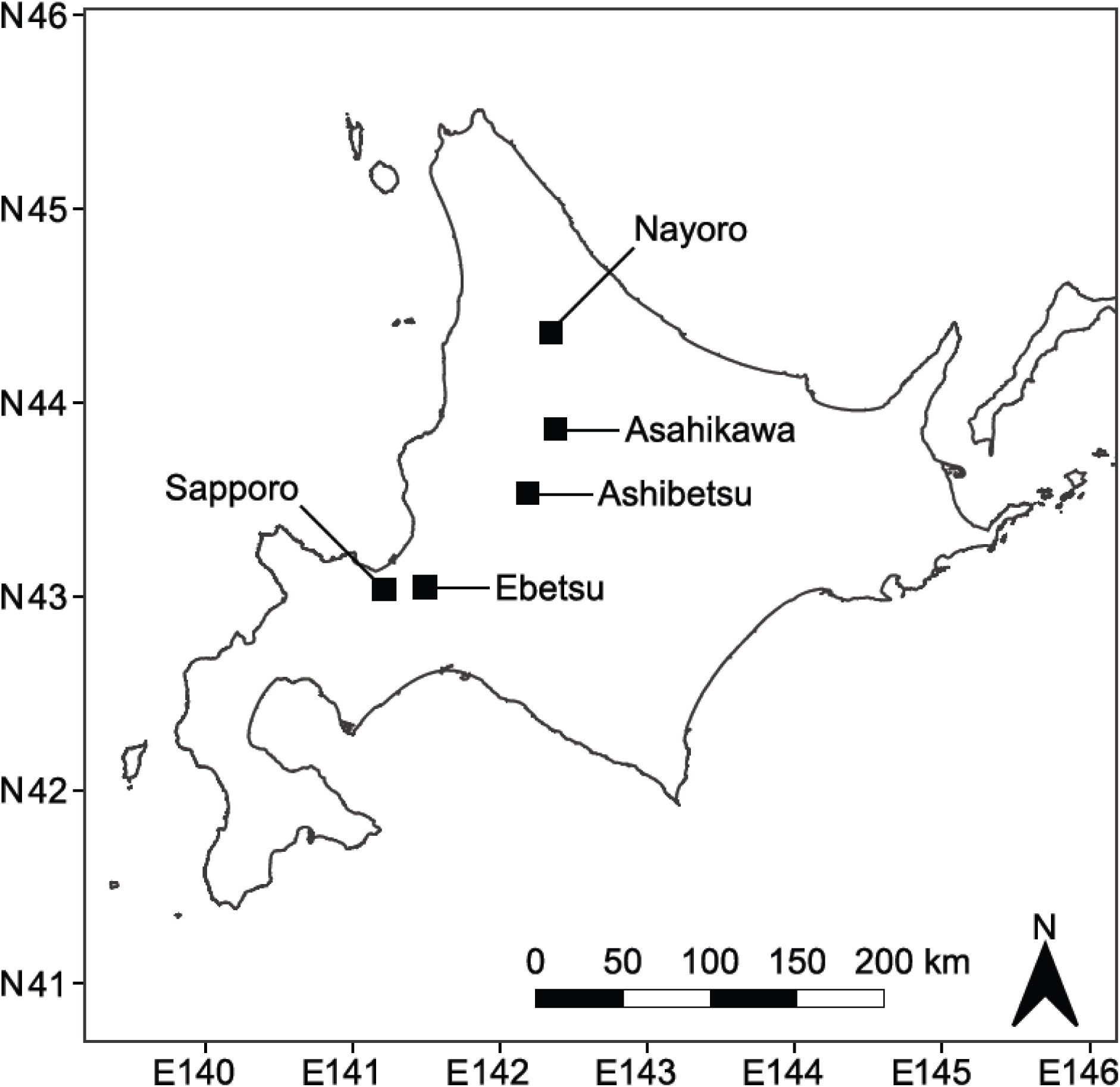
Study area in Hokkaido, northern Japan. Black squares indicate study regions.

**Table 1.** Habitat profile of study regions. Climate data indicate mean values for 2000–2020 at the nearest meteorological observation stations.

| Study sites | Study route length (km) | Average annual temperature (°C) | Total annual precipitation (mm) | Dominant tree species | Surrounding land use land cover |
| --- | --- | --- | --- | --- | --- |
| Sapporo | 5.5 | 9.2 | 1146.1 | *Acer pictum*,<br>*Tilia japonica* | Urban |
| Ebetsu | 12.5 | 7.3 | 965.0 | *Acer pictum*,<br>*Quercus crispula* | Urban,<br>Cropland |
| Ashibetsu | 6.9 | 7.4 | 1141.6 | *Abies sachalinensis*,<br>*Quercus crispula* | Deciduous broad-leaved forest |
| Asahikawa | 3.9 | 6.4 | 1081.7 | *Betula platyphylla*,<br>*Quercus crispula* | Paddy field |
| Nayoro | 3.5 | 5.8 | 1006.8 | *Betula platyphylla*,<br>*Quercus crispula* | Paddy field |

### Field Survey

Between April and May 2023, we surveyed the egg sacs of Ezo salamanders in ponds, puddles, and ditches along the survey routes in each region twice. We defined ponds and puddles as a single point covering the entire water area, and ditches as a single point every 10 m along their course. Because Ezo salamanders lay two egg sacs at a time, the number of egg sacs indicates the number of breeding populations (Sato 2013a). If an egg sac was found, its position and the number of egg sacs pairs were recorded using a handheld GPS device (Garmin eTrex 20x; Garmin Japan Co., Ltd.). If the number of egg sacs increased by the time of the second survey, the number recorded in the second survey was used. No significant decreases were observed during the second survey period. We randomly selected study sites that were more than 200 m apart and excluded the remaining sites from the survey to ensure spatial independence. During the second survey, additional study sites without egg sacs were randomly selected along the survey route, and their positions were recorded using handheld GPS devices. Absent study sites were randomly selected from the survey route at a distance of 200 m or more from study sites where egg sacs were found. The same number of absent and present study sites were selected. All study sites, including those with and without egg sacs, were set at 200 m or more away from each other. Moran’s I test showed no significant difference (*P*-values > 0.05) when the number of Ezo salamander egg sacs was used as the response variable for each survey area; therefore, the survey sites were considered independent.

### Landscape factors

We calculated the percentage of past and present forest areas within a 100 m buffer radius from the study sites to investigate the effects of past and present land use on the abundance or presence–absence of Ezo salamander egg sacs. This buffer size was chosen based on the movement distance of Ezo salamanders (Sato and Tsutsumi 2013). The percentage of past forest area was calculated by measuring forest areas in the 1960s aerial photographs published by the Geospatial Information Authority of Japan, Ministry of Land, Infrastructure, Transport, and Tourism. This period was chosen based on previous studies that have investigated the effects of past land use on salamanders (Cosentino and Brubaker 2018; Goldspiel et al. 2019). The percentage of the present forest area was calculated using the Japan High Resolution Land Use and Land Cover Map (Nov 2021 release, version 21.11, resolution 10 m), published by the Japan Aerospace Exploration Agency (https://www.eorc.jaxa.jp/ALOS/jp/dataset/lulc/lulc_v2111_j.htm). The map shows the average conditions from 2018 to 2020. Landscape factors were calculated using geographic information systems (QGIS 3.18; QGIS Development Team 2025).

### Local factors

We surveyed local factors at each study site. The following local factors were recorded: water temperature (℃), waterbody area (m²), water depth (cm), litter depth (cm), pH, electrical conductivity (mS m), dissolved oxygen (mg L), and waterbody type (lentic or lotic) to investigate the effects of local factors on the abundance or presence–absence of Ezo salamander egg sacs. The lentic water areas (ponds and puddles) were calculated as the product of length and width. Lotic water areas (ditches) were defined as having a length of 10 m and were calculated as the product of width and 10 m. Additionally, for local factors without water area or waterbody type, measurements were taken at three randomly selected points for ponds and puddles and at three points for ditches: the central point and points 5 m upstream and downstream. The average values were calculated. Water temperature, pH, and electrical conductivity were measured using a portable water quality meter (D-210PC-S, HORIBA Ltd.), and dissolved oxygen was measured using a waterproof dissolved oxygen meter (DO-1000PE, Custom Co. Ltd.). Waterbody type was defined as lentic for ponds and puddles and lotic for ditches as a categorical variable. Measurements of local factors were conducted during the second egg sac survey at all study sites.

### Data analysis

Principal component analysis was performed on the local factors, excluding the waterbody type, to reduce the number of explanatory variables and prevent overfitting of the model. A principal component analysis was performed for the following variables: water temperature, waterbody area, water depth, litter depth, pH, electrical conductivity, and dissolved oxygen. PC1 and PC2 were generated (Table 2). PC1 alone accounted for 32.2% of the total variation, whereas PC1 and PC2 together accounted for 55.6%. PC1 showed that higher positive values corresponded to greater litter depth and lower negative values corresponded to higher pH and dissolved oxygen. PC2 showed that higher positive values corresponded to wider water areas and higher water temperatures.

**Table 2.** Loadings of environmental factors and accumulative contributions of the three principal components. Bold style indicates a high factor loading (> |0.4|).

| Environmental factors | PC1 | PC2 |
| --- | --- | --- |
| Waterbody area (m<sup>2</sup>) | 0.341 | **0.542** |
| Water depth (cm) | 0.186 | 0.216 |
| Litter depth (cm) | **0.491** | 0.014 |
| Water temperature (°C) | 0.059 | **0.672** |
| pH | **-0.457** | 0.387 |
| Electrical conductivity (mS m) | -0.382 | 0.242 |
| Dissolved oxygen (mg L) | **-0.501** | 0.003 |
| Accumulative contribution (%) | 0.322 | 0.556 |

Generalized linear mixed models were constructed to investigate the effects of landscape and local factors on the abundance or presence–absence of Ezo salamander egg sacs. The response variable was the abundance or presence–absence of Ezo salamander egg sacs, and the explanatory variables were the percentage of past and present forest areas, PC1, PC2, and waterbody type, with region as the random effect. When the response variable was the abundance of egg sacs, a Poisson distribution was assumed, and the link function was set to log. When the response variable was the presence or absence of egg sacs, a binomial distribution was assumed, and the link function was set to logit. The categorical variable (waterbody type) was converted into a dummy variable, and all explanatory variables were standardized. The absolute values of the correlation coefficients between the explanatory variables were less than 0.7, indicating that collinearity between the explanatory variables had little effect on the results (Dormann et al. 2013). We constructed a group of models using all combinations of explanatory variables, averaged the models, and confirmed their significance (*P*-values < 0.05). To evaluate the effect size of each explanatory variable, 95% confidence intervals (CIs) of the standardized regression coefficients for each explanatory variable were calculated. All data analyses were performed using R version 4.4. (R Core Team 2025), and the R package “lme4” (Bates et al. 2025) and “MuMIn” (Bartoń 2025).

## Results

Ezo salamander egg sacs were found at 25 of the 50 study sites (both present and absent), with 245 pairs. At the sites where the salamanders were present, the average number of egg sacs per site (± SD) was 9.8 ± 9.6 pairs, ranging from a minimum of one pair to a maximum of 40 pairs.

The results of the generalized linear mixed models showed that the number of egg sacs was significantly associated with the percentage of past forest area, local factors (PC1), and waterbody type (Table 3). The standardized regression coefficients of the explanatory variables were high in the following order: local factors (PC1), waterbody type, and percentage of past forest area (Figure 2). The number of egg sacs was higher in areas with a higher percentage of past forest area, greater litter depth, and lower pH and dissolved oxygen in the lentic waters (ponds and puddles) (Figure 3). In addition, the presence or absence of egg sacs was significantly associated with local factors (PC1), with higher egg sac presence probabilities observed in areas with greater litter depth, lower pH, and lower dissolved oxygen (Table 3 and Figures 2 and 3).

**Fig. 2.**
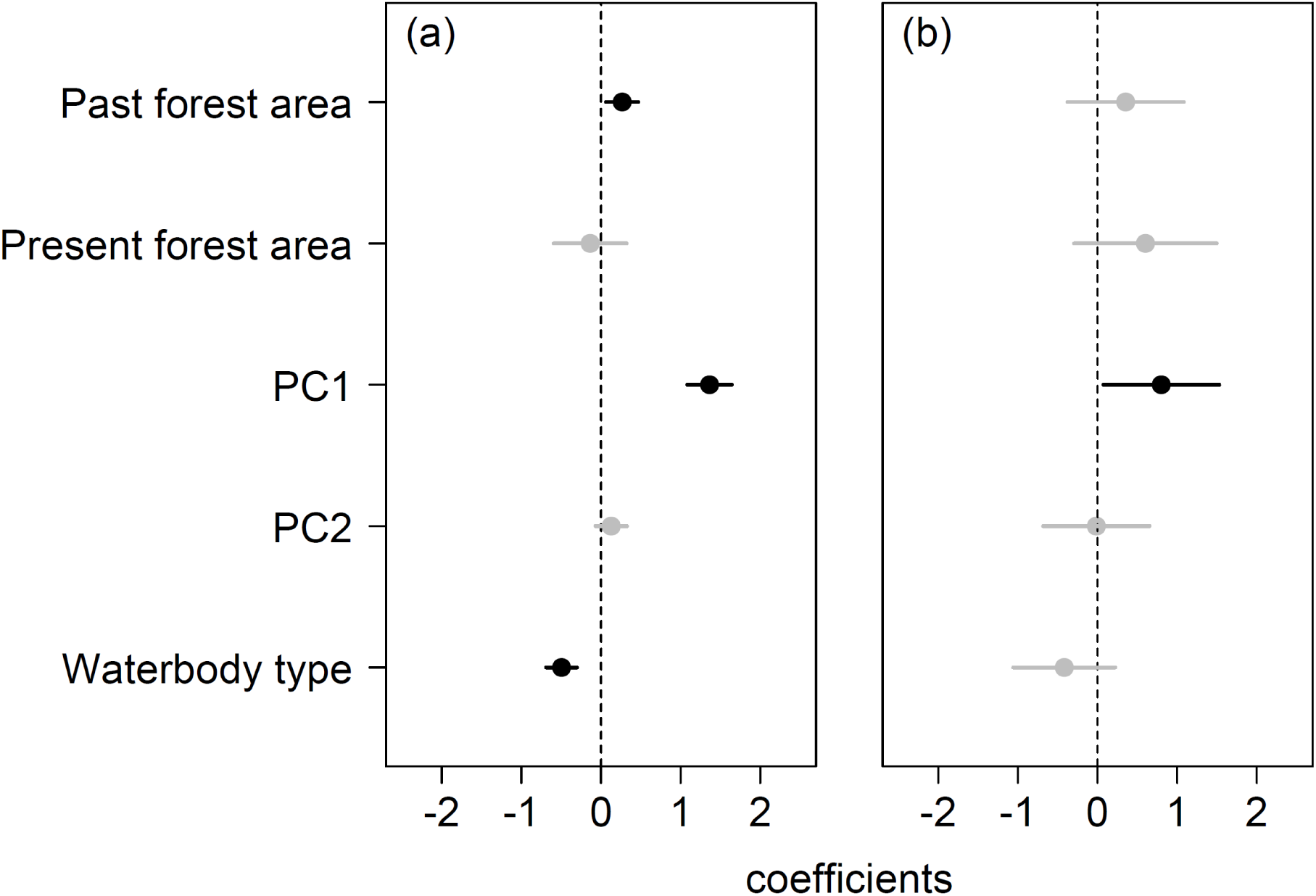
95% confidence intervals of standardized regression coefficients of each variable for (a) abundance, (b) presence–absence of Ezo salamander *Hynobius retardatus*. Black points and lines indicate significant effects, and gray points and lines indicate non-significant effects estimated using model averaging.

**Fig. 3.**
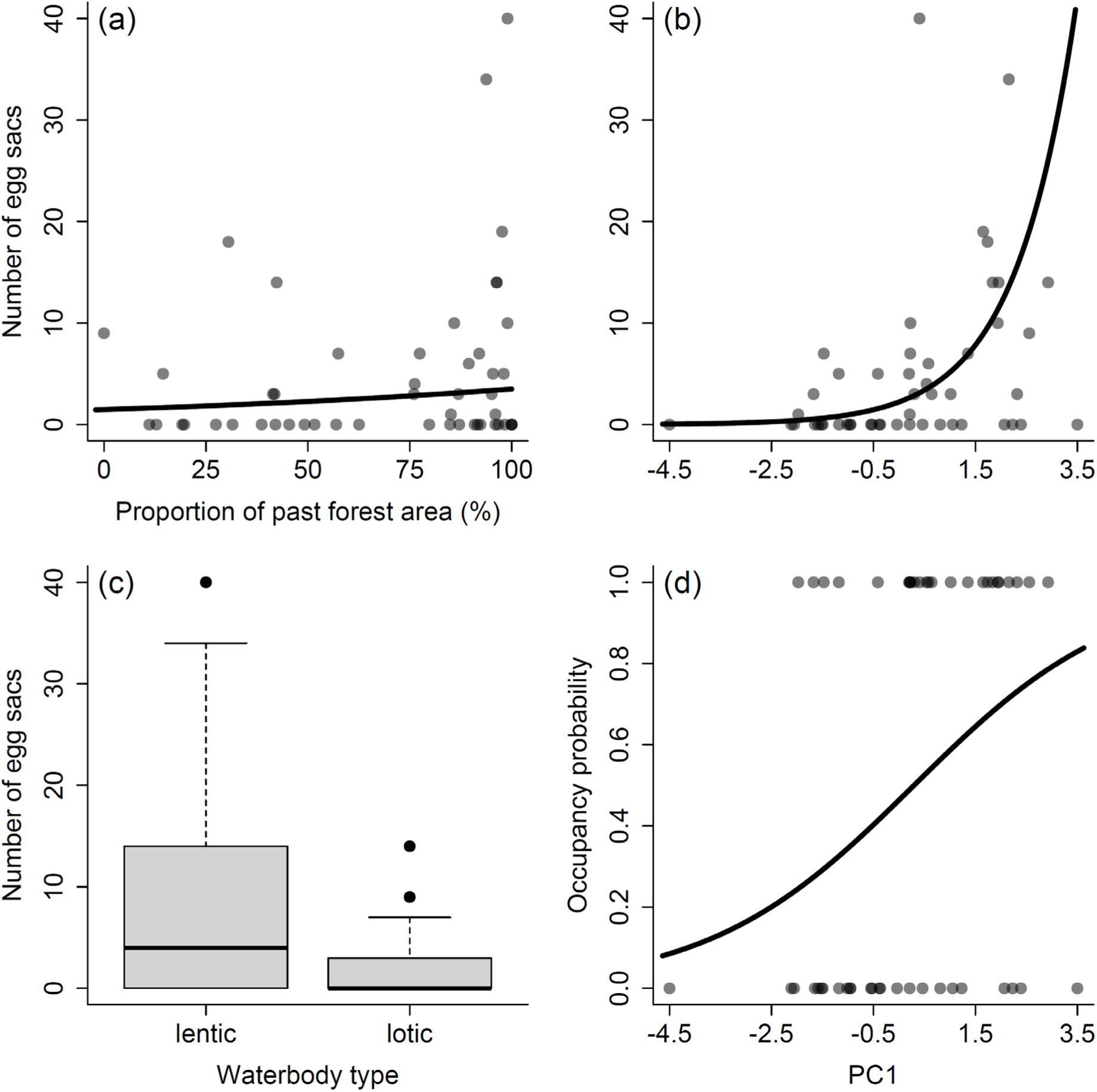
Relationship between (a) - (c) number of egg sacs (abundance) and (d) occupancy probability (presence–absence), and each variable. Black lines indicate significant effects estimated using model averaging.

**Table 3.** Parameter estimates, standard errors (SE), and *p*-values estimated using model averaging. Bold style indicates a significant parameter (*P*-values < 0.05).

| Responses | Variables | Estimates | SE | *P*-value |
| --- | --- | --- | --- | --- |
| Abundance | **Intercept** | **0.99** | **0.33** | **< 0.01** |
|  | **Past forest area** | **0.27** | **0.10** | **0.01** |
|  | Present forest area | -0.14 | 0.23 | 0.56 |
|  | **PC1** | **1.36** | **0.14** | **< 0.01** |
|  | PC2 | 0.13 | 0.10 | 0.21 |
|  | **Waterbody type** | **-0.50** | **0.10** | **< 0.01** |
| Presence–absence | Intercept | -0.04 | 0.31 | 0.91 |
|  | Past forest area | 0.35 | 0.36 | 0.34 |
|  | Present forest area | 0.60 | 0.44 | 0.19 |
|  | **PC1** | **0.80** | **0.36** | **0.03** |
|  | PC2 | -0.01 | 0.33 | 0.97 |
|  | Waterbody type | -0.42 | 0.32 | 0.20 |

## Discussion

The results of this study suggest that the effects of past land use on organism distribution are more pronounced when abundance data are used as the response variable than when presence–absence data are used. Previous studies on extinction debt have mostly focused on presence–absence data, with few comparisons made with abundance data (Hylander and Ehrlén 2013). We emphasize the importance of accounting for abundance in the context of extinction debt and suggest that focusing solely on the presence or absence may lead to an underestimation of extinction risk.

There are two possible reasons why extinction debt was more evident in abundance data than in presence–absence data. First, common species that are widely distributed tend to have high densities and variations in abundance, making their responses to environmental factors easier to detect. The Ezo salamander, the target species of this study, is listed on the Ministry of the Environment’s Red List, but is widely distributed in Hokkaido (Ministry of the Environment 2020; Tokuda 2022). In the present study, there were relatively high numbers of egg sacs per study site, with a maximum of 40 pairs; therefore, the relationship with environmental factors could be well explained using the abundance data. Similar to the results of this study, Cushman and McGarigal (2004) showed that presence–absence data for rare species and abundance data for common species better explain the relationship with present environmental factors. Second, presence–absence data may not fully reflect the environmental factors at a given study site, as they include sporadic and dispersed individuals. Previous studies have indicated that 50% of salamanders move within 41 m of their breeding sites (Rittenhouse and Semlitsch 2007). While many individuals settle near breeding sites, others move or disperse over distances ranging from hundreds of meters to several kilometers (Lowe et al. 2006; Semlitsch 2008). Therefore, individuals who moved long distances or were dispersed may have obscured the relationship between presence–absence data and environmental factors.

In the present study, the number of egg sacs was positively associated with the percentage of forested areas in the past. The red-backed salamander *Plethodon cinereus*, which inhabits North America, is scarce in post-agricultural forests due to reduced canopy cover and litter on the forest floor (Cosentino and Brubaker 2018). The results of our study corroborate these findings. While this study did not fully consider local factors on the forest floor, it is thought that past changes in the forest area led to a decline in habitat quality, resulting in a gradual decrease in abundance. In species such as the Ezo salamander, which has multiple life history stages in both terrestrial and aquatic environments, extinction does not occur immediately as long as the aquatic habitat remains unaffected (Hylander and Ehrlén 2013). Additionally, the extinction debt is more likely to occur when large-scale changes occur over a short period of time (Vellend et al. 2006; Kuussaari et al. 2009). In both Hokkaido, the focus of this study, and North America, the focus of previous studies, major land-use changes have occurred within only the past few hundred years (Ramankutty and Foley 1999; Hall et al. 2002; Yamanaka et al. 2015). As large-scale development has occurred over a short period, it is possible that an extinction debt was incurred. Therefore, a future study comparing findings from regions with different modification histories to those of Hokkaido and North America (e.g., regions where modifications occurred slowly over a long period) may provide a clearer understanding of the mechanisms of extinction debt.

In this study, the probability of egg sacs was positively associated with litter depth and negatively associated with pH and dissolved oxygen. As salamanders lay their eggs on fallen leaves and use them as hiding places for their larvae (Walls 1995; Chen et al. 2016), litter plays an important role in egg laying and larval growth. Furthermore, previous studies have shown that litter increases the abundance of microorganisms and their respiration, thereby reducing the dissolved oxygen levels (Rubbo et al. 2008). Decreases in pH and dissolved oxygen can cause acidification and hypoxia, which may adversely affect the survival of egg sacs and larvae (Kiesecker 1996; Mills and Barnhart 1999). Some salamander species are tolerant to changes in pH and dissolved oxygen, whereas others adapt through phenotypic plasticity (Sacerdote and King 2009; Fairman et al. 2013; Segev et al. 2019). Ezo salamanders also exhibit phenotypic plasticity, enlarging their gills under low-oxygen conditions (Iwami et al. 2007). Changes in water quality may not negatively affect the survival rates. Furthermore, decreased pH and dissolved oxygen levels may indirectly benefit Ezo salamanders by affecting their predators. The larvae of caddisflies, *Dytiscidae*, and fish prey on Ezo salamander egg sacs and larvae (Sato 2012, 2013b). In water areas with low pH and dissolved oxygen, there are fewer predators, such as aquatic insects; therefore, survival is possible (Janssen et al. 2018; Nieoczym et al. 2023; Malacarne et al. 2024). We found larvae of the caddisfly *Eubasilissa regina* at our study site in Ebetsu (Ohuchi personal observation). Therefore, it is thought that Ezo salamanders prefer water areas with a thick litter depth, low pH, and low levels of dissolved oxygen to increase the survival rate of their egg sacs and larvae.

We found that the number of egg sacs was higher in lentic water than in lotic water. Sato (1990) reported that 86.5% of 82 Ezo salamander laying sites were located in lentic waters with a flow velocity of less than 5 cm/s. These lentic waters not only prevent the outflow of egg sacs and larvae but may also reduce predation risks. Many of these water bodies are temporarily flooded during the spring and summer (Midori et al. 2017). They may become water bodies with low predation risk for Ezo salamanders by fish and other predators. Additionally, Ezo salamander egg sacs are often found in the same locations as Ezo brown frog *Rana pirica* egg masses (Sato 1990). In this study, egg masses of Ezo brown frogs were also found in the same locations as in the lentic water areas (Tamada, personal observation). Ezo salamander larvae prey on Ezo brown frog larvae, and when the two species coexist, the salamander larvae develop larger jaws (Kishida et al. 2006; Takatsu and Kishida 2013). Ezo-brown frog larvae are thought to play an important role as a food source. Therefore, there may be many Ezo salamander egg sacs in lentic waters where Ezo brown frog larvae are abundant.

In this study, local factors (PC1) were found to be the most effective explanatory variables when abundance was used as the response variable, followed by waterbody type and percentage of past forest area. This is because the number of egg sacs is strongly influenced by short-term direct factors, such as the physical and chemical conditions of the laying sites and the structure of the waterbody, rather than by long-term land use history. The drivers influencing adult occurrence and reproduction (eggs or larvae) of amphibians often differ; for example, forest area affects the probability of adult emergence of *Rana latastei* but does not affect reproduction probabilities(Lo Parrino et al. 2025). As this study used egg sacs as an indicator of the number of breeding populations of the Ezo salamander, which spawns in water bodies but inhabits forests as adults, the effect size of local factors may have been larger than that of the forest environment (Sato 2013a; Tokuda 2022). Second, the extinction debt may have been repaid over time. The study regions are located in areas connected to continuous forests and are not isolated. Therefore, the effects of past land use may have been offset by migration from these forests. Indeed, the number of Ezo salamander egg sacs is known to be higher in areas closer to continuous forests (Fuyuki et al. 2014).

Based on the results of this study, it is necessary to prioritize the conservation and improvement of physical environments and water quality in water areas when implementing habitat conservation or restoration for Ezo salamanders, while also considering the extinction lag caused by land-use changes. When devising conservation management plans that consider extinction lag, focusing on abundance data rather than presence–absence data may enable a more accurate assessment of extinction risk. Presence–absence data record, whether a species exists in a specific location. Compared to abundance data, presence–absence data are easier to collect and are therefore often used in citizen monitoring and wide-area surveys (Higa et al. 2015; Altwegg and Nichols 2019). However, caution is required when applying these data to extinct debt. Researchers and land managers should consider land-use strategies based on appropriate evaluation indicators such as abundance data.

## Acknowledgments

We are grateful to the members of Chodai Co., Ltd. for their assistance with the field survey and their valuable comments.

## Author Contributions

YT formulated this idea and developed the methodology. YT, NO, and KS conducted fieldwork. YT performed statistical analyses and wrote the manuscript. All authors contributed to drafting and approval of the final manuscript for publication.

## Ethics declarations

## Funding

None to declare.

### Ethical approval

Not applicable.

### Consent to participate

Not applicable.

### Consent for publication

Not applicable.

### Availability of data and material

The datasets used and/or analyzed in the current study are available from the corresponding author upon reasonable request.

### Code availability

The code used in this study is available from the corresponding author upon request.

## References

Altwegg R, Nichols JD (2019) Occupancy models for citizen-science data. Methods Ecol Evol 10:8–21. 10.1111/2041-210X.13090

Bartoń K (2025) Package ‘MuMIn.’ https://cran.r-project.org/web/packages/MuMIn/MuMIn.pdf. Accessed 1 Aug 2025

Bates D, Maechler M, Bolker B, Walker S, Christensen RHB, Singmann H, Dai B, Scheipl F, Grothendieck G, Green P, Fox J, Bauer A, Krivitsky PN, Tanaka E, Jagan M (2025) Package ‘lme4.’ https://cran.r-project.org/web/packages/lme4/lme4.pdf. Accessed 1 Aug 2025

Best ML, Welsh HH (2014) The trophic role of a forest salamander: impacts on invertebrates, leaf litter retention, and the humification process. Ecosphere 5:1–19. 10.1890/ES13-00302.1

Bryan BA, Gao L, Ye Y, Sun X, Connor JD, Crossman ND, Stafford-Smith M, Wu J, He C, Yu D, Liu Z, Li A, Huang Q, Ren H, Deng X, Zheng H, Niu J, Han G, Hou X (2018) China’s response to a national land-system sustainability emergency. Nature 559:193–204. 10.1038/s41586-018-0280-2

Chen C, Yang J, Wu Y, Fan Z, Lu W, Chen S, Yu L (2016) The breeding ecology of a critically endangered salamander, Hynobius amjiensis (Caudata: Hynobiidae), endemic to Eastern China. Asian Herpetol Res 7:53–58. 10.16373/j.cnki.ahr.150050

Cordier JM, Aguilar R, Lescano JN, Leynaud GC, Bonino A, Miloch D, Loyola R, Nori J (2021) A global assessment of amphibian and reptile responses to land-use changes. Biol Conserv 253:108863. 10.1016/j.biocon.2020.108863

Cosentino BJ, Brubaker KM (2018) Effects of land use legacies and habitat fragmentation on salamander abundance. Landsc Ecol 33:1573–1584. 10.1007/s10980-018-0686-0

Cushman SA, McGarigal K (2004) Patterns in the species-environment relationship depend on both scale and choice of response variables. Oikos 105:117–124. 10.1111/j.0030-1299.2004.12524.x

Davic RD, Welsh HH (2004) On the ecological roles of salamanders. Annu Rev Ecol Evol Syst 35:405–434. 10.1146/annurev.ecolsys.35.112202.130116

Dormann CF, Elith J, Bacher S, Buchmann C, Carl G, Carré G, Marquéz JRG, Gruber B, Lafourcade B, Leitão PJ, Münkemüller T, Mcclean C, Osborne PE, Reineking B, Schröder B, Skidmore AK, Zurell D, Lautenbach S (2013) Collinearity: A review of methods to deal with it and a simulation study evaluating their performance. Ecography 36:27–46. 10.1111/j.1600-0587.2012.07348.x

Exposito-Alonso M, Booker TR, Czech L, Gillespie L, Hateley S, Kyriazis CC, Lang PLM, Leventhal L, Nogues-Bravo D, Pagowski V, Ruffley M, Spence JP, Toro Arana SE, Weiß CL, Zess E (2022) Genetic diversity loss in the Anthropocene. Science 377:1431–1435. 10.1126/science.abn5642

Fairman CM, Bailey LL, Chambers RM, Russell TM, Funk WC (2013) Species-Specific Effects of Acidity on Pond Occupancy in Ambystoma Salamanders. J Herpetol 47:346–353. 10.1670/12-019

Figueiredo L, Krauss J, Steffan-Dewenter I, Sarmento Cabral J (2019) Understanding extinction debts: spatio–temporal scales, mechanisms and a roadmap for future research. Ecography 42:1973–1990. 10.1111/ecog.04740

Fuyuki A, Yamaura Y, Nakajima Y, Ishiyama N, Akasaka T, Nakamura F (2014) Pond area and distance from continuous forests affect amphibian egg distributions in urban green spaces: A case study in Sapporo, Japan. Urban For Urban Green 13:397–402. 10.1016/j.ufug.2013.11.003

Goldspiel HB, Cohen JB, McGee GG, Gibbs JP (2019) Forest land-use history affects outcomes of habitat augmentation for amphibian conservation. Glob Ecol Conserv 19: e00686. 10.1016/J.GECCO.2019.E00686

Grimm NB, Faeth SH, Golubiewski NE, Redman CL, Wu J, Bai X, Briggs JM (2008) Global Change and the Ecology of Cities. Science 319:756–760. 10.1126/science.1150195

Guirguis J, Goodyear LEB, Finn C, Johnson J V., Pincheira-Donoso D (2023) Risk of extinction increases towards higher elevations across the world’s amphibians. Global Ecology and Biogeography 32:1952–1963. 10.1111/geb.13746

Hall B, Motzkin G, Foster DR, Syfert M, Burk J (2002) Three hundred years of forest and land-use change in Massachusetts, USA. J Biogeogr 29:1319–1335. 10.1046/J.1365-2699.2002.00790.X

Hanski I (2011) Habitat loss, the dynamics of biodiversity, and a perspective on conservation. Ambio 40:248–255. 10.1007/s13280-011-0147-3

Hanski I, Ovaskainen O (2002) Extinction debt at extinction threshold. Conservation Biology 16:666–673. 10.1046/J.1523-1739.2002.00342.X

Herpetological Society of Japan (2025) Standard Japanese Names of Amphibians and Reptiles of Japan (ver. 2025-4-28). https://herpetology.jp/wamei/ (in Japanese). Accessed 1 Aug 2025

Higa M, Yamaura Y, Koizumi I, Yabuhara Y, Senzaki M, Ono S (2015) Mapping large-scale bird distributions using occupancy models and citizen data with spatially biased sampling effort. Divers Distrib 21:46–54. 10.1111/DDI.12255

Hylander K, Ehrlén J (2013) The mechanisms causing extinction debts. Trends Ecol Evol 28:341–346. 10.1016/J.TREE.2013.01.010

Iwami T, Kishida O, Nishimura K (2007) Direct and indirect induction of a compensatory phenotype that alleviates the costs of an inducible defense. PLoS One 2: e1084. 10.1371/JOURNAL.PONE.0001084

Jackson ST, Sax DF (2010) Balancing biodiversity in a changing environment: extinction debt, immigration credit and species turnover. Trends Ecol Evol 25:153–160. 10.1016/j.tree.2009.10.001

Janssen A, Hunger H, Konold W, Pufal G, Staab M (2018) Simple pond restoration measures increase dragonfly (Insecta: Odonata) diversity. Biodivers Conserv 27:2311–2328. 10.1007/S10531-018-1539-5/FIGURES/5

Kiesecker J (1996) pH-mediated predator-prey interactions between Ambystoma Tigrinum and Pseudacris Triseriata. Ecological Applications 6:1325–1331. 10.2307/2269610

Kishida O, Mizuta Y, Nishimura K (2006) Reciprocalocal phenotypic plasticity in a predator-prey interaction between larval amphibians. Ecology 87:1599–1604. 10.1890/0012-9658(2006)87

Koshika K, Nigi T (1998) The current state and challenges of forest practices tor urban forests − A case study of the Nopporo Forest Park, Hokkaido. Jpn J For Plann 41–49. 10.20659/jjfp.30.0_41 (in Japanese)

Kuussaari M, Bommarco R, Heikkinen RK, Helm A, Krauss J, Lindborg R, Öckinger E, Pärtel M, Pino J, Rodà F, Stefanescu C, Teder T, Zobel M, Steffan-Dewenter I (2009) Extinction debt: a challenge for biodiversity conservation. Trends Ecol Evol 24:564–571. 10.1016/j.tree.2009.04.011

Lo Parrino E, Ficetola GF, Devin M, Manenti R, Falaschi M (2025) Integrating adult occurrence and reproduction data to identify conservation measures for amphibians. Conservation Biology 39: e14343. 10.1111/COBI.14343

Lowe WH, Likens GE, Cosentino BJ (2006) Self-organisation in streams: the relationship between movement behaviour and body condition in a headwater salamander. Freshw Biol 51:2052–2062. 10.1111/J.1365-2427.2006.01635.X

Malacarne TJ, Machado NR, Moretto Y (2024) Influence of land use on the structure and functional diversity of aquatic insects in neotropical streams. Hydrobiologia 851:265–280. 10.1007/s10750-023-05207-5

Midori T, Kuwahara T, Yamashiki N (2017) Retardation of larval development in the salamander Hynobius retardatus in a permanent pond with abundant spring water. Limnology 18:287–299. 10.1007/S10201-016-0506-7

Mills NE, Barnhart MC (1999) Effects of hypoxia on embryonic development in two Ambystoma and two Rana species. Physiological and Biochemical Zoology 72:179–188. 10.1086/316657

Ministry of the Environment J (2020) Publication of the Ministry of the Environment’s Red List 2020. In: https://www.env.go.jp/press/107905.html (in Japanese) Accessed 1 Aug 2025

Navarro LM, Pereira HM (2012) Rewilding Abandoned Landscapes in Europe. Ecosystems 15:900–912. 10.1007/S10021-012-9558-7/FIGURES/5

Newbold T, Hudson LN, Hill SLL, Contu S, Lysenko I, Senior RA, Börger L, Bennett DJ, Choimes A, Collen B, Day J, De Palma A, Díaz S, Echeverria-Londoño S, Edgar MJ, Feldman A, Garon M, Harrison MLK, Alhusseini T, Ingram DJ, Itescu Y, Kattge J, Kemp V, Kirkpatrick L, Kleyer M, Correia DLP, Martin CD, Meiri S, Novosolov M, Pan Y, Phillips HRP, Purves DW, Robinson A, Simpson J, Tuck SL, Weiher E, White HJ, Ewers RM, MacE GM, Scharlemann JPW, Purvis A (2015) Global effects of land use on local terrestrial biodiversity. Nature 520:45–50. 10.1038/nature14324

Nieoczym M, Stryjecki R, Buczyński P, Płaska W, Kloskowski J (2023) Differential abundance, composition and mesohabitat use by aquatic macroinvertebrate taxa in ponds with and without fish. Aquat Sci 85:25. 10.1007/s00027-022-00922-y

Pounds JA, Bustamante MR, Coloma LA, Consuegra JA, Fogden MPL, Foster PN, La Marca E, Masters KL, Merino-Viteri A, Puschendorf R, Ron SR, Sánchez-Azofeifa GA, Still CJ, Young BE (2006) Widespread amphibian extinctions from epidemic disease driven by global warming. Nature 439:161–167. 10.1038/nature04246

QGIS Development Team (2025) QGIS geographic information system. QGIS a free and open source geographic information system. https://qgis.org/en/site/. Accessed 1 Aug 2025

Queiroz C, Beilin R, Folke C, Lindborg R (2014) Farmland abandonment: threat or opportunity for biodiversity conservation? A global review. Front Ecol Environ 12:288–296. 10.1890/120348

R Core Team (2025) R: A language and environment for statistical computing. R Foundation for Statistical Computing. http://www.R-project.org. Accessed 1 Aug 2025

Ramankutty N, Foley JA (1999) Estimating historical changes in land cover:North American croplands from 1850 to 1992. Global Ecology and Biogeography 8:381–396. 10.1046/J.1365-2699.1999.00141.X

Re:wild, Synchronicity Earth, IUCN SSC Amphibian Specialist Group (2023) State of the World’s Amphibians: The Second Global Amphibian Assessment. Re:wild, Texas, USA

Rittenhouse TAG, Semlitsch RD (2007) Distribution of amphibians in terrestrial habitat surrounding wetlands. Wetlands 27:153–161

Rubbo MJ, Belden LK, Kiesecker JM (2008) Differential responses of aquatic consumers to variations in leaf-litter inputs. Hydrobiologia 605:37–44. 10.1007/s10750-008-9298-z

Sacerdote AB, King RB (2009) Dissolved oxygen requirements for hatching success of two ambystomatid salamanders in restored ephemeral ponds. Wetlands 29:1202–1213. 10.1672/08-235.1

Sato T (1990) Temperature and velocity of water at breeding sites of Hynobius retardatus. Japanese Journal of Herpetology 13:131–135

Sato T (2012) Ecology and conservation of two salamander species distribution in Hokkaido. Hokkaido herpetological society research report 1:47–48 (in Japanese)

Sato T (2013a) Let’s learn about salamander ecology. In: Sato T, Matsui M (eds) Salamanders of Hokkaido. Eco Network, Sapporo, pp 5–15 (in Japanese)

Sato T (2013b) Natural enemies and diseases. In: Sato T, Matsui M (eds) Salamanders of Hokkaido. Eco Network, Sapporo, pp 96–98 (in Japanese)

Sato T, Tsutsumi K (2013) Case study at Wakaba Forest, Obihiro City. In: Sato T, Matsui M (eds) Salamanders of Hokkaido. Eco Network, Sapporo, pp 155–167 (in Japanese)

Segev O, Pezaro N, Rovelli V, Rybak O, Templeton AR, Blaustein L (2019) Phenotypic plasticity and local adaptations to dissolved oxygen in larvae fire salamander (Salamandra infraimmaculata). Oecologia 190:737–746. 10.1007/S00442-019-04446-5/FIGURES/4

Semlitsch RD (2008) Differentiating migration and dispersal processes for pond-breeding amphibians. J Wildl Manage 72:260–267. 10.2193/2007-082

Takatsu K, Kishida O (2013) An offensive predator phenotype selects for an amplified defensive phenotype in its prey. Evol Ecol 27:1–11. 10.1007/S10682-012-9572-4

Thornton DH, Branch LC, Sunquist ME (2011) The influence of landscape, patch, and within-patch factors on species presence and abundance: A review of focal patch studies. Landsc Ecol 26:7–18. 10.1007/s10980-010-9549-z

Tilman D, Fargione J, Wolff B, D’Antonio C, Dobson A, Howarth R, Schindler D, Schlesinger WH, Simberloff D, Swackhamer D (2001) Forecasting agriculturally driven global environmental change. Science (1979) 292:281–284. 10.1126/SCIENCE.1057544

Tilman D, May RM, Lehman CL, Nowak MA (1994) Habitat destruction and the extinction debt. Nature 371:65–66. 10.1038/371065a0

Tokuda T (2022) Compact picture guide to Hokkaido reptiles & amphibians. The Hokkaido Shimbun Press, Sapporo (in Japanese)

Tran D Van, Terui S, Nomoto K, Nishikawa K (2021) Ecological niche differentiation of two salamanders (Caudata: Hynobiidae) from Hokkaido Island, Japan. Ecol Res 36:281–292. 10.1111/1440-1703.12191

Ueta T, Sato T (2013) Age composition. In: Sato T, Matsui M (eds) Salamanders of Hokkaido. Eco Network (in Japanese), Sapporo, pp 93–95

Vellend M, Verheyen K, Jacquemyn H, Kolb A, Van Calster H, Peterken G, Hermy M (2006) Extinction debt of forest plants persists for more than a century following habitat fragmentation. Ecology 87:542–548. 10.1890/05-1182

Walls SC (1995) Differential vulnerability to predation and refuge use in competing larval salamanders. Oecologia 101:86–93

Winkler K, Fuchs R, Rounsevell M, Herold M (2021) Global land use changes are four times greater than previously estimated. Nat Commun 12:1–10. 10.1038/s41467-021-22702-2

Yamanaka S, Akasaka T, Yamaura Y, Kaneko M, Nakamura F (2015) Time-lagged responses of indicator taxa to temporal landscape changes in agricultural landscapes. Ecol Indic 48:593–598. 10.1016/j.ecolind.2014.08.024

Zhang K, Dang H, Tan S, Wang Z, Zhang Q (2010) Vegetation community and soil characteristics of abandoned agricultural land and pine plantation in the Qinling Mountains, China. For Ecol Manage 259:2036–2047. 10.1016/J.FORECO.2010.02.014

